# Molecular characterization of latent GDF8 reveals mechanisms of activation

**DOI:** 10.1101/155614

**Authors:** Ryan G. Walker, Jason C. McCoy, Magdalena Czepnik, Melanie J. Mills, Adam Hagg, Kelly L. Walton, Thomas Cotton, Marko Hyvönen, Richard T. Lee, Paul Gregorevic, Craig A. Harrison, Thomas B. Thompson

**Author notes:** co-corresponding authors Correspondence: University of Cincinnati 231 Albert Sabin Way ML 0524 Cincinnati, OH 45267, Monash University 26 Innovation Walk Clayton VIC 3800. equal contribution (Ryan G. Walker, Jason C. McCoy).

## Abstract

Growth/differentiation factor 8 (GDF8) or myostatin negatively regulates muscle mass. GDF8 is held in a latent state through interactions with its N-terminal prodomain, much like TGF-β. Using a combination of small angle X-ray scattering and mutagenesis, we characterized the interactions of GDF8 with its prodomain. Our results show that the prodomain:GDF8 complex can exist in a fully latent state and an activated or ‘triggered’ state where the prodomain remains in complex with the mature domain. However, these states are not reversible, indicating the latent GDF8 is ‘spring-loaded’. Structural analysis shows that the prodomain:GDF8 complex adopts an ‘open’ configuration, distinct from the latency state of TGF-β and more similar to the ‘open’ state of Activin A and BMP9 (non-latent complexes). We determined that GDF8 maintains similar features for latency, including the alpha-1 helix and fastener elements, and identified a series of mutations in the prodomain of GDF8 that alleviate latency, including I56E, which does not require activation by the protease Tolloid. *In vivo,* active GDF8 variants were potent negative regulators of muscle mass, compared to wild-type GDF8. Collectively, these results help characterize the latency and activation mechanisms of GDF8.

## Introduction

One of the most thoroughly described negative regulators of skeletal muscle mass is the TGF-β superfamily ligand growth/differentiation factor 8 (GDF8), also known as myostatin (1, 2). Genetic disruption of *GDF8* results in substantial skeletal muscle growth (1-7). Further, a significant increase in muscle fiber size is also observed when adult animals are treated with agents that bioneutralize GDF8 (reviewed in (8)). As such, targeted inhibition of GDF8 is currently being pursued for the treatment of skeletal muscle-related disorders and associated symptoms (9, 10).

GDF8, like numerous TGF-β family members, is a disulfide-linked dimer that is synthesized as a precursor protein which requires cleavage by a furin-like protease to yield a N-terminal prodomain and a C-terminal mature, signaling domain (11). Interestingly, for a number of TGF-β ligands the role of the prodomain extends beyond ligand maturation and folding support (12, 13), remaining non-covalently associated with the mature ligand following secretion in either a low-affinity, non-inhibitory or high-affinity, inhibitory fashion (reviewed in (14)). For example, the prodomains of TGF-β1, TGF-β2, TGF-β3, GDF11, and GDF8 hold the mature ligand in a latent or inactive state mediated by a non-covalent, yet high affinity, ligand-specific interaction (11, 15-18) whereas mature Activin A and BMP9 remain associated with, but are not inhibited by, their prodomain (19, 20). Activation of TGF-β1 and TGF-β3 requires covalent interactions with the extracellular matrix and cellular contractile forces to release the mature ligand (21-23). In fact, resolution of the latent TGF-β1 crystal structure provided a molecular explanation for how latency is exerted by the prodomain via a coordinated interaction between the N-terminal alpha helix (alpha1), latency lasso, and fastener of the prodomain with type I and type II receptor epitopes of the mature domain (23). On the other hand, GDF8 activation requires a second cleavage event within the prodomain via proteases from the BMP1/Tolloid (TLD) family of metalloproteases (24). However, the molecular and structural details of the GDF8 latent state have yet to be determined.

Based on sequence conservation and prior biochemical data describing the N-terminal portion of the GDF8 prodomain (15), it is plausible that the molecular interactions and overall structure of the GDF8 latent complex may be similar to that of TGF-β1. However, the prodomains of a number of TGF-β family members share similar sequence conservation, yet they do not regulate the mature ligand in the same fashion and also exhibit significant structural diversity (19, 20). Therefore, while one might expect that GDF8 and TGF-β1 would share certain elements for how the prodomain binds and confers latency, it is possible that significant structural and molecular differences in these interactions occur as they exhibit profoundly different mechanisms of activation. However, this comparison is hindered by a lack of understanding of the GDF8 latent complex at the molecular level.

In this study, we utilized small angle X-ray scattering (SAXS) and mutagenesis to characterize the GDF8 latent complex. Interestingly, SAXS analysis reveals that the GDF8 latent complex adopts a more ‘open’ conformation, similar to the overall structure of the BMP9 and Activin A prodomain complexes, which are not latent. The ‘open’ conformation of the GDF8 latent complex is in stark contrast to the ‘closed’ conformation adopted by the TGF-β1 latent complex. Furthermore, we identify key residues in the GDF8 prodomain that are responsible for promoting latency indicating that GDF8 and TGF-β1 share similar features for latency including a latency lasso. We further show that certain mutations in the prodomain of GDF8 can reduce latency, producing a more active ligand both *in vitro* and *in vivo*. Overall, our data provide insight toward the molecular mechanisms of GDF8 latency and activation.

## Results

### Prodomain-GDF8 can exist in a latent and active complex

Initial characterization in adult mice showed that GDF8 is secreted into the systemic circulation as a latent protein complex that requires activation to trigger GDF8 signaling (16). While the biological mechanism for activation remained unknown, it was shown that a GDF8-specific signal derived from the serum of a wild-type mouse, but not a *gdf8*^-/-^ mouse, could be detected following exposure to acidic conditions, referred to here as “acid-activation” (16). The premise for acid-activation stemmed from a similar observation that was made during the characterization of TGF-β, which is similarly regulated by its prodomain, (25, 26) and provided the initial basis that latent GDF8 and latent TGF-β are likely to be very similar in terms of activation and prodomain release.

While a molecular basis to describe how acid-activation alleviates ligand latency remains unknown, it is thought that the acidic-conditions simply dissociate the prodomain from the mature domain, thereby freeing the ligand from inhibition (16). However, our initial attempts to purify the mature domain from the prodomain after acid-activation using an affinity column to the high-affinity antagonist, follistatin, failed, even though the complex exhibited significant activity. This observation suggested that perhaps the prodomain remained bound to the mature domain, but was not in a fully inhibitory state. To extend these initial observations, we isolated the mammalian-derived latent proGDF8 complex (GDF8^L^) and compared its signaling activity to both the acid-activated state (GDF8^AA^) and to the mature, unbound GDF8 (GDF8^apo^) using a SMAD3-responsive (CAGA)_12_ luciferase reporter HEK293 cell line (27-31). As expected and consistent with our previous report, GDF8^apo^ readily signaled with a calculated half-maximal effective concentration (EC_50_) of 0.72 nM (31), whereas media containing GDF8^L^ did not readily signal and required nearly 10,000-fold more protein to achieve a similar response as compared to GDF8^apo^ (Figure 1a). In contrast, acid-activation of media containing GDF8^L^ at pH 2 to generate GDF8^AA^ resulted in a significant gain in activity compared to non-acid activated latent GDF8 (Figure 1a). Interestingly, the calculated EC_50_ for GDF8^AA^ (5.7 nM) still did not reach the EC_50_ of GDF8^apo^, suggesting that under these conditions we were unable to observe the full signaling potential of mature GDF8. Since GDF8^apo^ is stable and stored in 10 mM HCl, we do not expect this difference in activity to be caused by subjecting GDF8^L^ to extreme conditions. We next evaluated the activation of proGDF8 as a function of pH by subjecting GDF8^L^ to various pH ranges (pH 2-10) for 1 hour, followed by neutralization and (CAGA)_12_ activation (Figure 1b). We determined that at the concentration tested (40 nM), the level of activation increases with a decrease in pH (Figure 1b), however substantial activation was observed throughout the pH range examined. The shape of the titration experiment indicated that multiple ionizable groups could be involved in the latency mechanism or that shifts in the pH cause disruption in the structure of the prodomain that effects its ability to inhibit GDF8.

**Figure 1.**
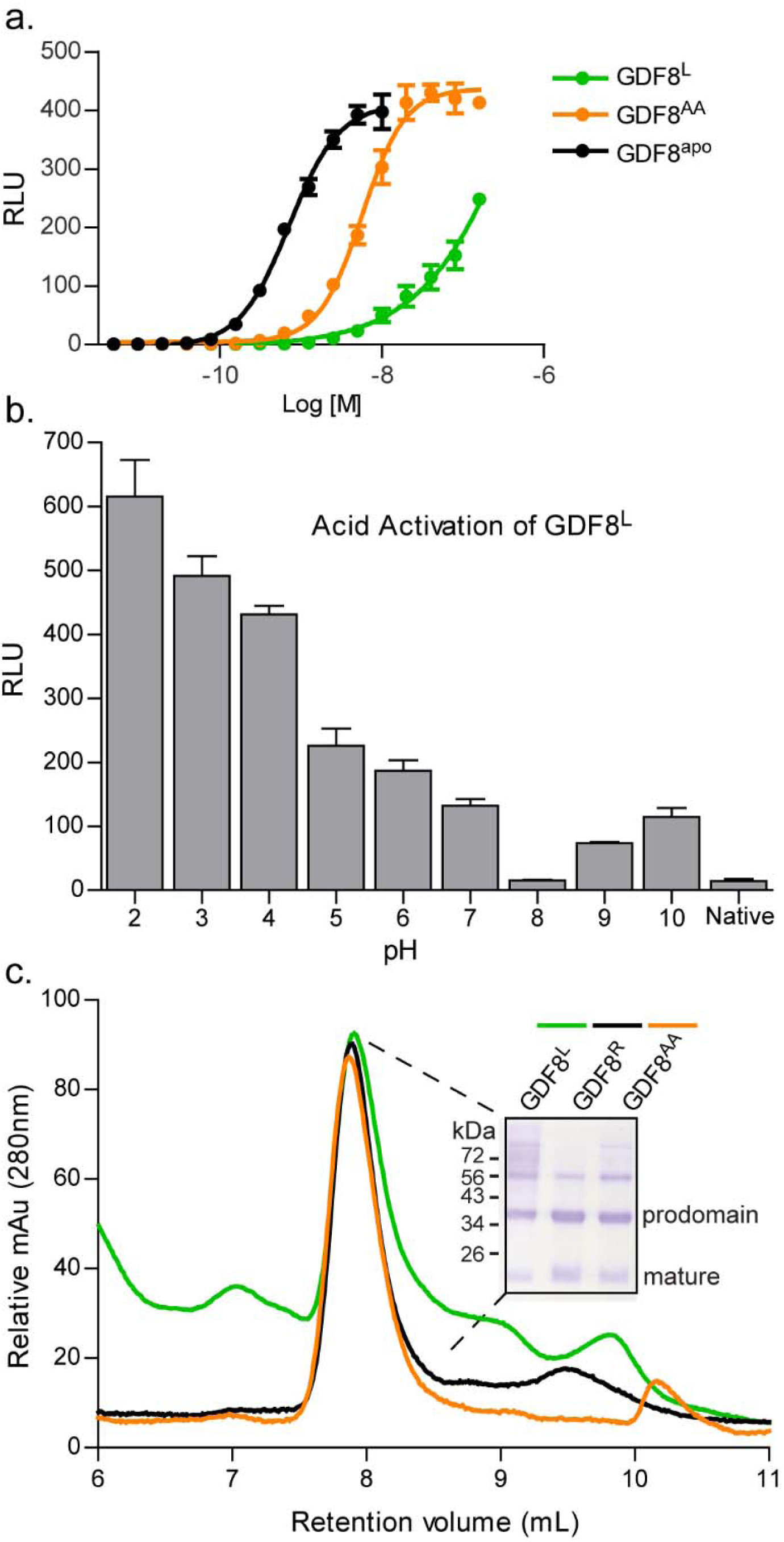
Activity and analysis of the latent GDF8 prodomain complex. **a)** Reporter assay using HEK293 (CAGA)_12_ cells treated with a titration of latent GDF8 prodomain complex (GDF8^L^), acid activated (GDF8^AA^), and mature (GDF8^apo^) ligand to detect activation. Experiments were performed at least twice with each data point measured in triplicate. Shown is a representative experiment. Data were fit by non-linear regression to a variable slope to determine the EC_50._ **b)** Activity measurement in HEK293 (CAGA)_12_ cells of GDF8^L^ (40nM). Error is shown as the mean ± standard error of the mean (SEM) **c)** SEC analysis of GDF8^L^, reformed (GDF8^R^), and GDF8^AA^ to detect for the presence of the prodomain:ligand complex. Inlet shows an SDS-PAGE of the peak.

Given that titration of the GDF8 prodomain against mature GDF8 results in potent ligand inhibition (15), we hypothesized that we were unable to recover the full signal from acid activated latent GDF8 due to the possibility that the prodomain may still be able to provide some level of antagonism, through a non-covalent interaction, despite being acid activated. To test this hypothesis, we subjected GDF8^L^ and GDF8^AA^ to size exclusion chromatography (SEC) followed by SDS-PAGE/Coomassie staining. In addition, we combined isolated GDF8 prodomain with GDF8^apo^ and applied the mixture to SEC. We determined that all three variations of GDF8 complexes had similar retention volumes with both components (co-eluting the prodomain and GDF8^apo^), indicative of complex formation (Figure 1c). This result supports the idea that during the acid activation, the prodomain can re-associate with the mature domain and partially inhibit signaling. However, since the GDF8^AA^ has significant activity, it also suggests that the latent interaction between the prodomain and mature domain is not completely reversible. Nevertheless, our finding that following exposure to acidic conditions, mature GDF8 remains associated with the prodomain, but in an active state, provided the opportunity for further comparison to latent GDF8.

### SAXS analysis reveals conformational differences between GDF8 and other TGFβ prodomain-ligand complexes

While the aforementioned data suggests that the latent and acid-activated forms of GDF8 could adopt different molecular states, limited structural information is available for the prodomain:GDF8 complex. Given that high-resolution structural information for prodomain:ligand complexes of latent TGF-β1 (23) and non-latent ligands, BMP9 (19) and Activin A (20) have demonstrated a series of configurations that range in compactness, we next wanted to determine how the prodomain:GDF8 complex compared to these other prodomain:ligand complexes. Therefore, we used the solution-based technique small angle X-ray scattering (SAXS) to analyze the purified prodomain:GDF8 complexes, including GDF8^L^, GDF8^R^, and GDF8^AA^ (Figure 2 and Table 1). Samples were well behaved in solution and did not show evidence of interparticle repulsion or aggregation over multiple protein concentrations (Figure 2a and Table 1). From the Gunier analysis, we determined that GDF8^L^ has a lower *R*_g_ than GDF8^AA^, 41.1 ± 0.85 *versus* 46.8 ± 0.86 Å (Table 1), respectively, which suggests that acid activation of GDF8^L^ altered the overall conformation of the complex. This is further supported by the appearance of a more ‘featured’ pairwise distribution plot (*P*(*r*)) for the GDF8^AA^ complex compared to GDF8^L^ complex (Figure 2b). Additionally, we determined that the GDF8^R^ complex had a similar scattering profile, pairwise distribution curve, and associated SAXS derived values as the GDF8^L^ complex (Figure 2a,b and Table 1).

**Table 1:** Experimentally determined parameters from SAXS analysis of GDF8 prodomain complexes.

| Sample | Concentration (mg/mL) | $I(0)$ (cm<sup>-1</sup>) | $R_g$ (Å) | | $D_{max}$ (Å) | Volume (Å<sup>3</sup>) | Mass (kDa) |
| --- | --- | --- | --- | --- | --- | --- | --- |
|  |  |  | Gunier | Real Space |  |  |  |
| Native | 3 | 4,400 | 41.7 | 39.3 | 136 | 310,000 | 90.0 |
|  | 2 | 2,400 | 40.5 | 38.7 | 133 | 300,000 | 76.0 |
| Acid Activated | 1.3 | 1,600 | 47.6 | 43.0 | 148 | 340,000 | 77.0 |
|  | 1.15 | 1,700 | 47.0 | 41.6 | 147 | 350,000 | 70.0 |
|  | 1 | 1,500 | 45.9 | 40.0 | 134 | 340,000 | 59.0 |
| Reformed | 1.6 | 190 | 40.0 | 39.0 | 131 | 270,000 | 89.0 |
| *Theoretical<sup>a</sup>* |  |  |  |  |  |  |  |
| proTGF-β1 | NA | NA | 28.5 | NA | 92 | NA | 82.4 |
| proBMP9 | NA | NA | 37.8 | NA | 137 | NA | 90.2 |
| proActivin A | NA | NA | 31.6 | NA | 106 | NA | 90.1 |
| proGDF8 | NA | NA | 34.9 | NA | 120 | NA | 80.1 |

<sup>a</sup>Values derived from deposited PDB coordinates for each prodomain:ligand complex

**Figure 2.**
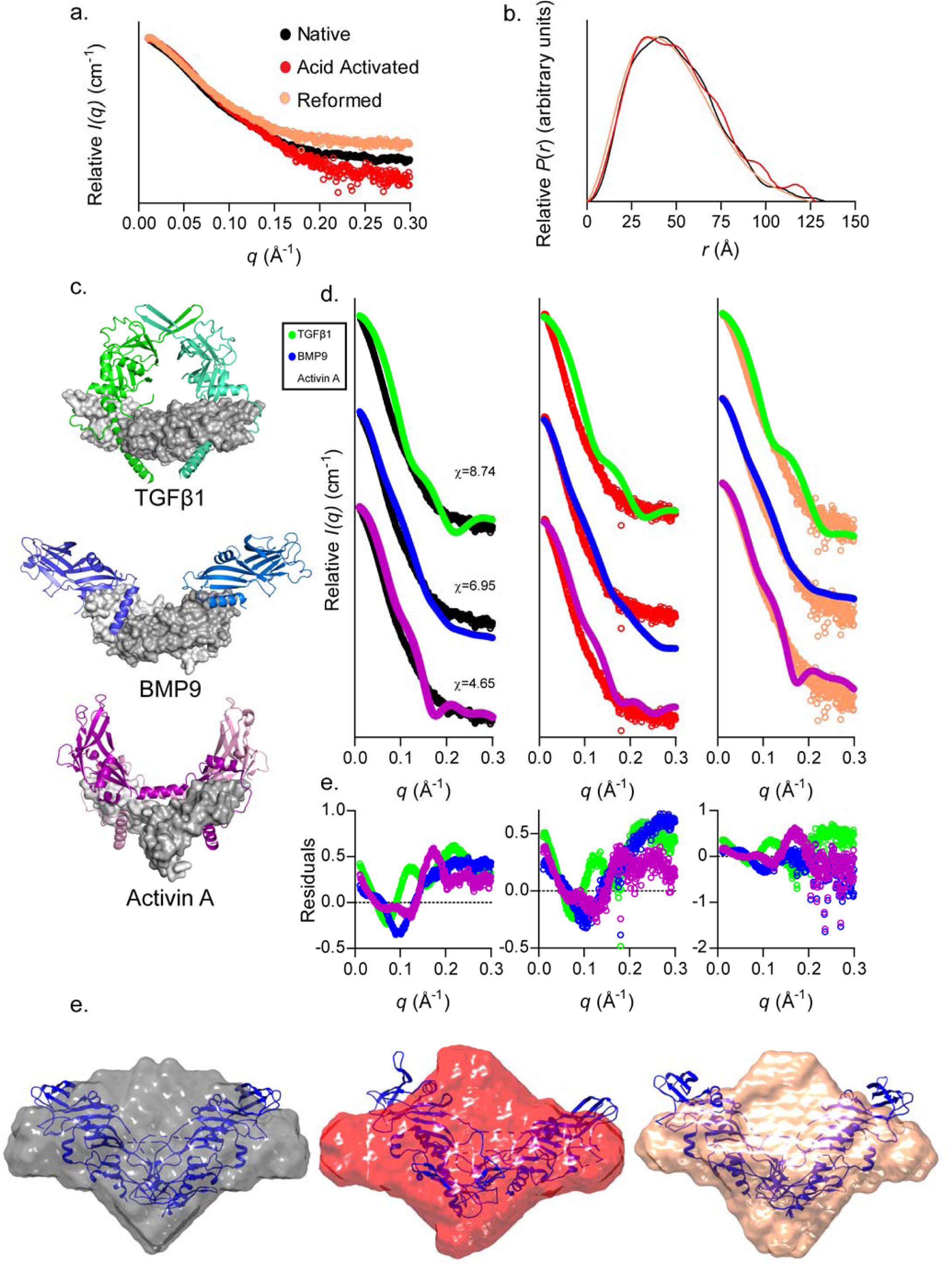
SAXS analysis of latent, acid activated, and reformed GDF8 prodomain complex. **a)** SAXS scattering profile showing the intensity distribution and (**b**) the pairwise distribution function for the various GDF8 prodomain complexes. **c)** and **d)** The known structures of the various prodomain:ligand complexes that were used to generate the theoretical scattering profile curves for comparison to our experimentally determined profiles of the various GDF8 prodomain complexes. The chi (X) value, determined by FoXS (32), for each comparison is shown adjacent to each scattering profile. Note that the latent TGF-β1 structure exemplifies a ‘closed’ conformation unlike the non-latent, but prodomain:ligand associated, BMP9 and Activin A structures are in an ‘open’ conformation. **e)** *Ab initio* SAXS envelope (DAMFILT model) of the native (black), acid activated (red), and reformed (pink) GDF8 prodomain complexes with the latent GDF8 prodomain complex structure superimposed (blue) (43).

To determine if the GDF8^L^ complex adopted a similar conformation to the other known prodomain:ligand structures, we compared our experimental scattering profile to the theoretical scattering profile using FoXS (Figure 2c; (32, 33)). We first compared the experimental profile of GDF8^L^ to the theoretical profiles based on the prodomain:ligand structures of TGF-β1, BMP9, and Activin A. This analysis showed that the overall structure of the GDF8^L^ complex did not show substantial similarity to any structure as indicated by the calculated chi values (Figure 2d), which is also consistent with a larger *R*_g_ value than the other prodomain:ligand structures. Nevertheless, the most similarity was found with Activin A (χ=4.65), which has an ‘open’ conformation, whereas the least similarity was found with TGF-β1 (χ=8.74; Figure 2c). Interestingly, both the GDF8^AA^ and GDF8^R^ complexes were more similar to Activin A (χ=1.96 and χ=1.43, respectively) while still a poor fit with BMP9 (χ=3.13 and χ=3.48, respectively) and TGFβ-1 (χ=3.35 and χ=3.04, respectively), suggesting that GDF8^AA^ and GDF8^R^ are also likely in an ‘open’ conformation and that there are likely additional differences in these complexes compared to the GDF8^L^ complex. To extend these observations, we calculated the SAXS-derived *ab initio* molecular envelopes for each state. The overall shape of the envelopes for each state further supported our initial observation that structural differences likely exist between the activity states (Figure 2e). However, there are poorly defined regions within the envelopes, which may be the result of structural flexibility inherent to the GDF8 prodomain complexes.

### Specific mutations within the prodomain enhance GDF8 activity

Our SAXS analysis revealed that the GDF8^L^ complex likely adopts a different overall conformation compared to TGF-β1. Despite this, GDF8 and TGFβ-1 share high sequence conservation in the N-terminal alpha-1 helix, latency lasso, alpha-2 helix, and fastener regions (Figure 3a). Thus, we hypothesized that these regions could interact with the mature GDF8 ligand and are important for forming the non-covalent interactions required for latency, such that removing these interactions might generate a more active GDF8 ligand (i.e. remove latency). One might also expect that disruption of these interactions might disrupt folding, as observed for TGF-β1 (34). Therefore, to test our hypothesis, we utilized the TGF-β1 structure as a guide to systematically mutate specific residues in regions of the GDF8 prodomain and compared their activity to wild-type GDF8^L^. For this evaluation we developed a robust cell-based (CAGA)_12_ luciferase-reporter assay where we could assess the variants through transient transfection. Our first goal was to determine which TLD family protease member (e.g. BMP1/mTLD, tolloid-like 1 (TLL1) or tolloid-like 2 (TLL2)) yielded the most optimal activation of wild-type GDF8^L^ (24, 35). Using a similar assay format as previously described (31), we compared the activity of wild-type GDF8 following transient co-transfection of wild-type GDF8, furin, and either BMP1, TLL1 or TLL2 using HEK293 (CAGA)_12_ luciferase cells (Figure 3b). As predicted, we observed little to no signal when the TLDs were not included in the assay, indicating that little to no basal TLD is present and incapable of activating GDF8^L^ (Figure 3b). However, when cells were co-transfected with DNA from one of the three TLDs, we observed a dose-dependent increase in signal with increasing concentrations of wild-type GDF8 DNA (Figure 3b). As predicted, we observed differences in the fold activation of wild-type GDF8 when co-transfected with the various TLDs, where the highest activation resulted from TLL2 (TLL2>TLL1>BMP1; Figure 3b). Although this result is consistent with previous reports (24, 35), suggesting that differences in the magnitude of activation by TLDs, we cannot rule out the possibility this increase in activity is due to differences in TLD protein expression levels or differential regulation of TLD maturation needed for activation (36-38). Regardless, since TLL2 was the most effective activator of wild-type GDF8 with increases ranging from 20 to 60-fold activation, it was used in the remaining assays unless otherwise noted.

**Figure 3.**
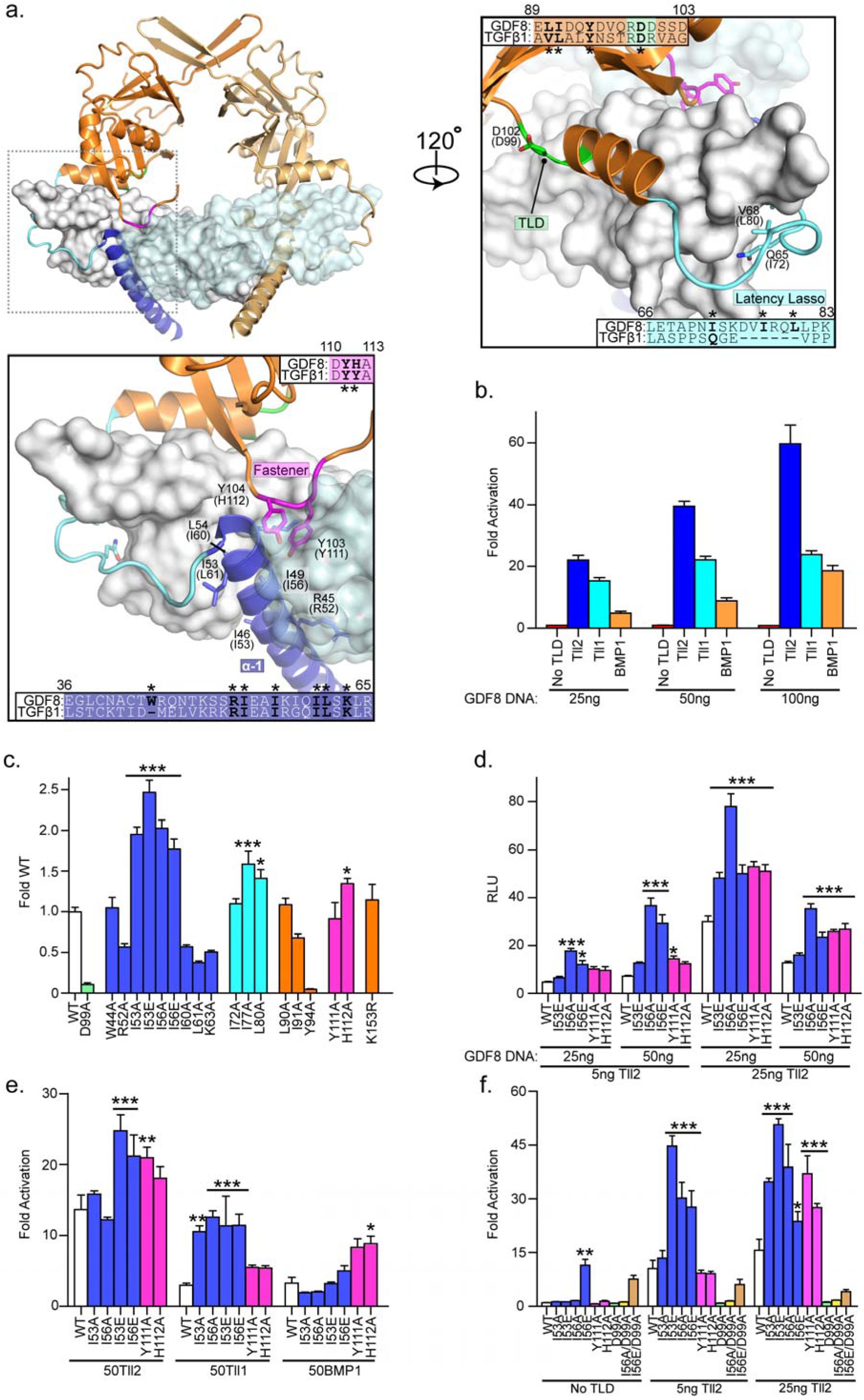
Mutations within the GDF8 prodomain and activation by Tolloid. **a)** The structure of TGF-β1 showing the important inhibitory elements: alpha-1 (blue), latency lasso (cyan) and the fastener (magenta), sticks representing mutants being made corresponding to GDF8 prodomain, bolded and marked with an asterisk within the sequence alignment. Numbering of residues utilizes the full-length propeptide including the signal sequence. **b)** Tolloid fold activation over background of WT GDF8 prodomain complex within a transfection based luciferase assay. 25ng of the indicated tolloid, TLL1, TLL2 or BMP1 was used in the presence of 50ng Furin. **c)** Screen of all mutants within the GDF8 prodomain in with 25ng TLL2, 25ng GDF8 construct, and 50ng furin DNA. Bar colors represent the region within the prodomain, blue (alpha-1), cyan (latency lasso), orange (prodomain), magenta (fastener) and reported as fold over wild type (WT) GDF8. **d)** Using pSF-IRES vector, mutants were tested within a transfection luciferase assay and normalized using Renilla luminescence reported as Relative Light Units (RLU). **e)** Using 50ng of each tolloid DNA and 25ng of the respective GDF8 mutant DNA within the co-transfected with an IRES vector for normalization, fold activation over no tolloid reported. **f)** A co-transfection of 25ng IRES vector and 25ng pRK5 expression vector for each construct at 0, 5, 25ng TLL2 was transfected and activity of each mutant examined and reported as relative light units (RLU). All mentioned experiments were performed at least twice were individual points were measured in triplicate. Error is shown as mean ± SEM. Bar graphs were compared using two-way ANOVA with Bonferroni correction (*P ≤ 0.05, **P ≤ 0.01, and ***P ≤ 0.001).

The panel of mutations is shown in Figure 3c and is categorized based on the anticipated location in the prodomain. Within these regions, we primarily focused our attention on mutation of hydrophobic residues, since hydrophobic interactions commonly drive known inhibitory interactions within the TGF-β family (reviewed in (14)). For example, GDF8 maintains a number of hydrophobic resides that are predicted to align to one side of the alpha-1 helix, similar to the register of TGF-β1. Of particular interest, we identified two hydrophobic residues in the alpha-1 helix, I53 and I56, which showed more than 2-fold higher activity compared to wild-type GDF8 when mutated to either an alanine (I53A, I56A) or glutamate (I53E, I56E; Figure 3c). Additionally, mutation of residues outside of the alpha-1 helix, I77A within the latency lasso, and H112A within the fastener showed an increase in activity when compared to wild-type, whereas the mutants generated in the alpha-2 helix did not show any significant gain in activity compared to wild-type (Figure 3c). In contrast, Y94A resulted in little to no activity. As a control, we tested the activity of D99A, which has previously been shown to eliminate activation by TLD (24). As expected, introduction of D99A abolished activity, supporting that the assay is specific to the plasmid carrying the GDF8 gene. In addition, we tested the K153R mutant, which was previously shown to enhance furin processing but not influence activation by TLD. K153R had similar activity as wild-type, indicating that TLD processing is optimal (39).

In order to validate these observations and perform a more rigorous cross-comparison between WT GDF8 and these mutants, we inserted the ligand DNA into the pSF-CMV-FMDV-Rluc vector, which allowed us to normalize our data for transfection efficiency. We focused on the I53A/E and I56A/E mutants within the alpha-1, as well as the Y111A and H112A mutants within the fastener region due to their apparent importance when examining the structure of TGF-β1. This approach was used because previous efforts to detect the secreted ligand in the conditioned medium in this assay format were unsuccessful, likely due to protein levels below the limit of detection. We determined that all mutants retain significantly higher activity than WT GDF8 in a dose-dependent fashion with respect to titration of ligand DNA and TLL2 DNA (Figure 3d). Given that activation of WT GDF8 is differentially regulated by the various TLDs, we tested whether or not our mutants retained higher activity when activated by the other TLDs, TLL1 and BMP1. Overall, our results indicated that our mutants were more active than WT GDF8, though there were a few differences in the activation across the various TLDs (Figure 3e). Except for the I53A and I56A variants of the Ile mutations co-transfected with TLL2, all mutants showed enhanced activity in the presence of either TLL2 or TLL1, while only the Y111A and H112A mutants showed enhanced activity when co-transfected with BMP1 (Figure 3e). These results were unexpected and likely suggest that the enhanced activity of our mutants may occur because of multiple mechanisms, such as whether or not TLD is still required for activation.

To determine if the enhanced ligand activity was dependent on TLD activity (i.e. TLD-dependent), we compared the activity of these mutants transfected with and without TLL2 (Figure 3f). Interestingly, of the mutants tested, the I56E mutant showed significant activity compared to WT GDF8 in the absence of TLL2 (Figure 3f), whereas the other mutants (I53A/E, Y111A, and H112A) required the presence of TLD for enhanced activity (Figure 3f). To confirm that GDF8 with the I56E mutation is not dependent on TLD, we generated the double mutant, I56E/D99A, which would eliminate the potential for TLL2 activation. Similar to I56E, transfection of the I56E/D99A mutant showed enhanced activity thus demonstrating that the I56E mutation results in non-latent and active GDF8 ligand (Figure 3f). However, we did observe that co-transfection of TLL2 further enhanced the activity of the I56E mutant suggesting that more activity from this mutant can still be gained, but not in the presence of D99A. Thus, I56E has activity without the requirement of TLD, but TLD can further potentiate I56E’s activity. The I56A mutant did not show the same TLL2 independence as I56E (Figure 3f) suggesting that introduction of the charged residue may destabilize the interaction between the prodomain and the mature ligand, perhaps by disrupting a hydrophobic pocket or core.

### GDF8 prodomain mutations exhibit reduced antagonism

As mentioned earlier, GDF8 mature ligand signaling can be antagonized by titrating increasing amounts of purified GDF8 prodomain. Therefore, we next wanted to determine if the prodomains with mutations we identified as activating had an altered capacity to inhibit the mature GDF8 ligand. To accomplish this, we produced and purified the GDF8 prodomain mutants in bacteria (Figure 4a) and determined their half-maximal inhibitory potential (IC_50_) against a constant concentration of mammalian derived, mature GDF8 (Figure 4b). To improve the production and solubility of the bacteria-derived GDF8 prodomain mutants, we mutated all four cysteines to serine (GDF8^4xCtoS^; *see Materials and Methods*). Using the SMAD3-responsive (CAGA)_12_ luciferase reporter HEK293 cell line described above, we determined the IC_50_ for several of the activating prodomain mutations (Figure 4b; Table 2). Results show that mutations in the fastener region, Y111A and H112A, had a similar IC_50_ to GDF8^4xCtoS^ whereas, mutations in the alpha 1 helix (I53A, I56A and I56E) were 3 to 4-fold less potent. However, the most dramatic effect was observed with I56E, which was ∼16-fold less potent than GDF8^4xCtoS^.

**Table 2:**
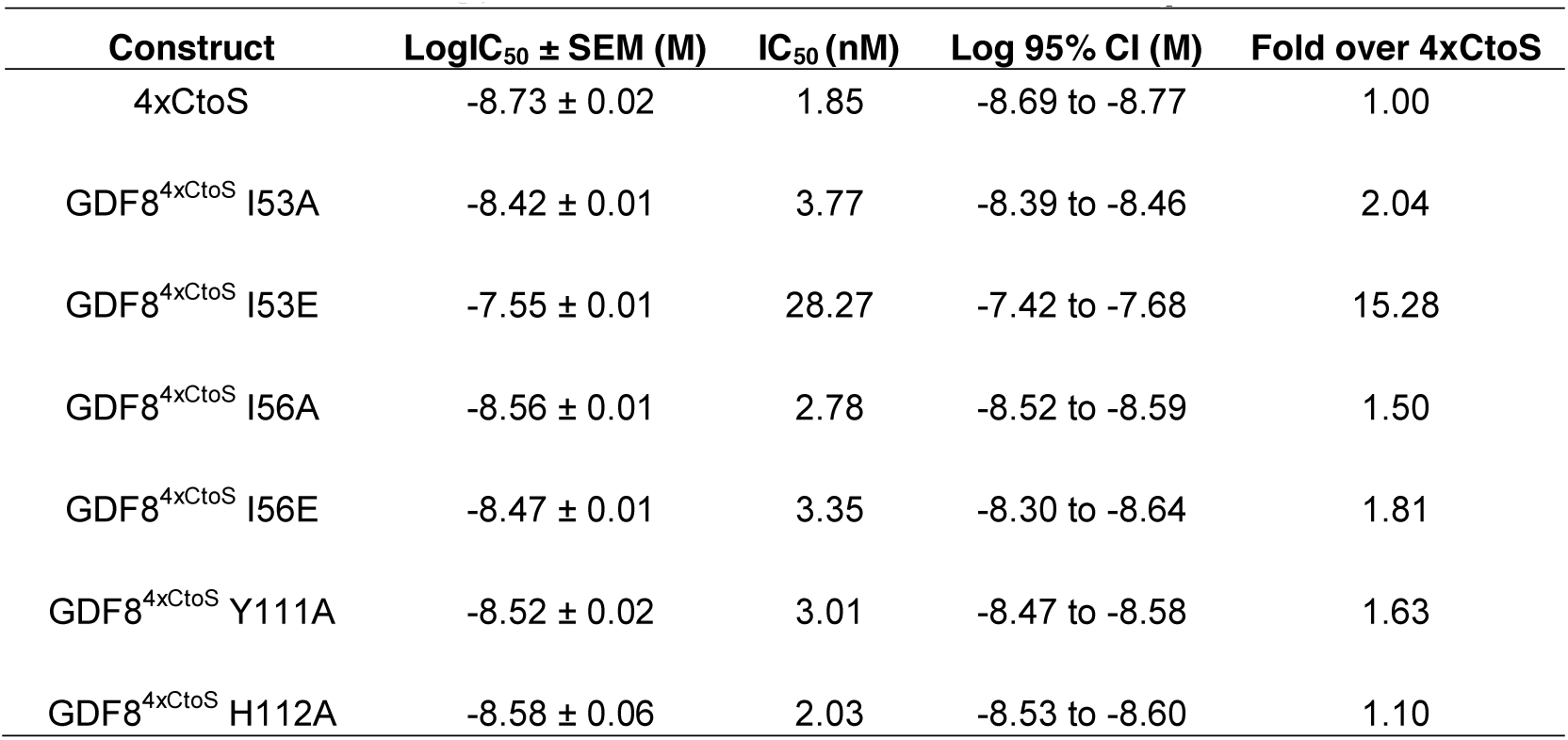
Calculated IC_50_ values for various mutant GDF8 prodomain constructs.

| Construct | LogIC<sub>50</sub> ± SEM (M) | IC<sub>50</sub> (nM) | Log 95% CI (M) | Fold over 4xCtoS |
| --- | --- | --- | --- | --- |
| 4xCtoS | -8.73 ± 0.02 | 1.85 | -8.69 to -8.77 | 1.00 |
| GDF8<sup>4xCtoS</sup> I53A | -8.42 ± 0.01 | 3.77 | -8.39 to -8.46 | 2.04 |
| GDF8<sup>4xCtoS</sup> I53E | -7.55 ± 0.01 | 28.27 | -7.42 to -7.68 | 15.28 |
| GDF8<sup>4xCtoS</sup> I56A | -8.56 ± 0.01 | 2.78 | -8.52 to -8.59 | 1.50 |
| GDF8<sup>4xCtoS</sup> I56E | -8.47 ± 0.01 | 3.35 | -8.30 to -8.64 | 1.81 |
| GDF8<sup>4xCtoS</sup> Y111A | -8.52 ± 0.02 | 3.01 | -8.47 to -8.58 | 1.63 |
| GDF8<sup>4xCtoS</sup> H112A | -8.58 ± 0.06 | 2.03 | -8.53 to -8.60 | 1.10 |

**Figure 4.**
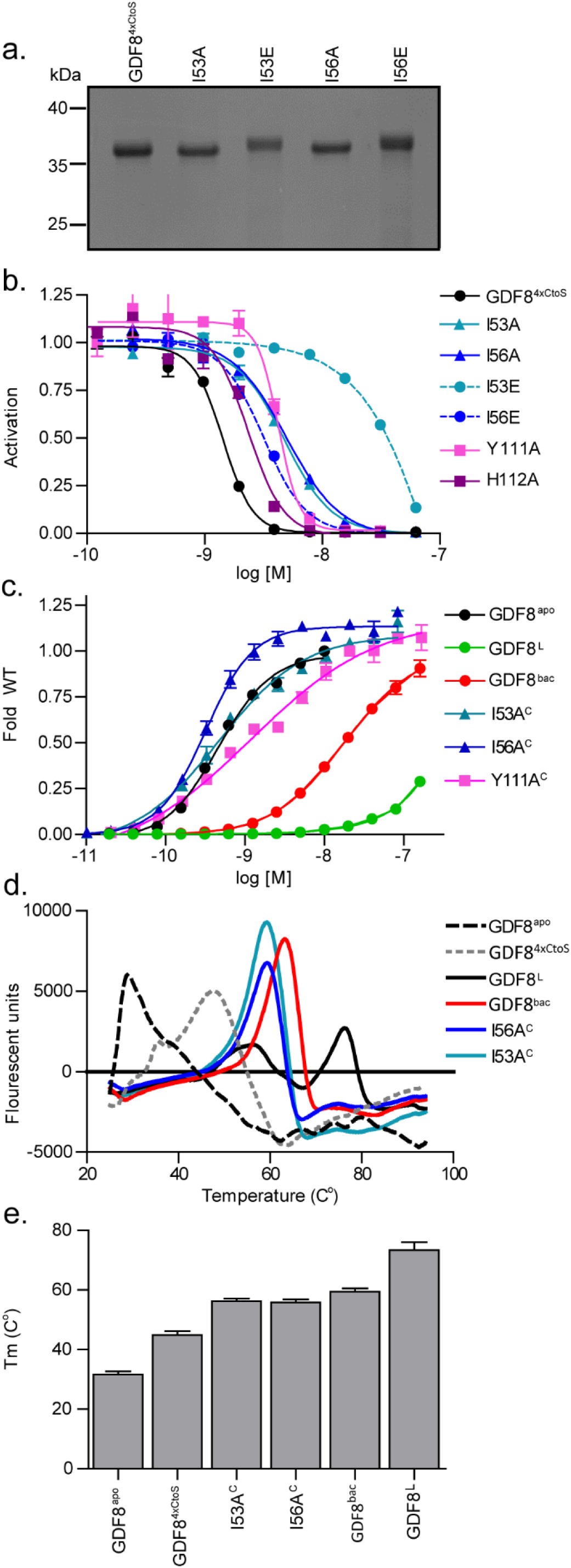
Bacterial-produced GDF8 prodomains. **a)** SDS gel of purified of GDF8^4xCtoS^ bacterially expressed prodomain mutants. **b)** Representative IC_50_ curve of serially diluted bacterial-expressed prodomains mixed with exogenous GDF8^apo^ and added to our HEK 293 (CAGA)_12_ cell line, activity reported as fold WT activity. **c)** EC_50_ values of reformed GDF8 complex with mutant prodomains. The protein complex was serially diluted using 1:2 dilutions to generate a fold WT curve. **e)** Representative derivative plot of melt curves generated via Thermal Shift and reported as fluorescent units. All mentioned experiments were performed at least twice were individual points were measured in duplicate. Error is shown as mean ± SEM.

### Reformed complexes using the GDF8 prodomain mutants are more active and exhibit decreased thermal stability

We next wanted to determine if we could reform the prodomain:ligand complex using the various mutant prodomain constructs and subsequently assess their signaling activity. Therefore, we combined the prodomain mutations with the mature ligand and isolated the complex by SEC. We calculated the EC_50_ of the complexes using the HEK293 (CAGA)_12_ luciferase-reporter cells and compared these results to GDF8^L^ complex, GDF8^4xCtoS^ complex, and GDF8^apo^ (Figure 4c; Table 3). We were unable to isolate a stable complex using the prodomain mutations of I53E and I56E, presumably due to a loss in affinity for the mature GDF8. Interestingly, all reformed mutant complexes showed significant activity with EC_50_ values similar to the mature GDF8 indicating that the prodomain:ligand inhibitory complex was less stable during the assay and could not function to inhibit GDF8 signaling. This is in contrast to the GDF8^4xCtoS^ complex, which had significantly less activity, but still had more activity than GDF8^L^ complex.

**Table 3:** Calculated EC_50_ values for various mutant GDF8 prodomain constructs.

| Construct | LogEC<sub>50</sub> ± SEM (M) | EC<sub>50</sub> (nM) | Log 95% CI (M) | Fold over 4xCtoS |
| --- | --- | --- | --- | --- |
| GDF8<sup>apo</sup> | -8.82 ± 0.03 | 1.53 | -8.75 to -8.88 | 1.00 |
| GDF8<sup>L</sup> | NC<sup>a</sup> | NC | NC | NC |
| GDF8<sup>4xCtoS</sup> | -7.93 ± 0.02 | 10.62 | -7.92 to -8.03 | 6.94 |
| GDF8<sup>4xCtoS</sup> I53A | -9.23 ± 0.03 | 0.59 | -8.90 to -9.57 | 0.39 |
| GDF8<sup>4xCtoS</sup> I56A | -9.53 ± 0.06 | 0.30 | -8.74 to -10.32 | 0.20 |
| GDF8<sup>4xCtoS</sup> Y111A | -8.98 ± 0.09 | 1.05 | -7.78 to -10.18 | 0.69 |

<sup>a</sup>NC=not calculable

To further determine if the enhanced activity shown by the mutants, specifically the alpha-1 mutants (I53, I56), may be explained in part by destabilization of the prodomain:mature ligand complex, we performed a thermal shift assay. In this assay the binding of the hydrophobic dye Rox dye was measured as a function of temperature (Figure 4d, e). We determined that the mammalian-derived GDF8^L^ complex had the highest T_m_ whereas the reformed GDF8^4xCtoS^ complex and GDF8 I53A and I56A mutant complexes showed a lower T_m_ suggestive of diminished stability or differences in the binding mode compared to the GDF8^L^ complex (Figure 4d, e). Both the GDF8^L^ and mutant complexes showed increased stability when compared to GDF8^apo^ and the unbound GDF8^4xCtoS^ prodomain indicating that the difference in T_m_ is not due to excess GDF8^apo^ ligand within the sample or as a result of dissociated, unbound prodomain (Figure 4d, e). In addition to the higher T_m_ maxima for the GDF8^L^ complex, we detected a second maxima at a lower temperature, not observed in the reformed complexes, suggesting that a complex destabilization event occurred for the GDF8^L^ complex (Figure 4d). Taken together, these data suggest that mutation of the residues within the alpha-1 helix alleviates GDF8 latency through disruption of the interaction between the prodomain and mature domain.

### GDF8 mutants enhance muscle atrophy compared to wild-type GDF8

Having demonstrated that mutation of specific residues within the GDF8 prodomain result in a more active or less latent ligand *in vitro* assays. Hence, we next wanted to determine if the enhanced activity would be recapitulated *in vivo* in a model of skeletal muscle atrophy. To test this, we generated AAV vectors encoding either WT GDF8 or the activating GDF8 mutants, I56E and H112A, and locally injected them in into the tibialis anterior (TA) muscles of 6-8-week-old male C57Bl/6 mice. Eight weeks post-AAV injections, WT GDF8, which is secreted in a latent form, induced a modest (∼7%) decrease in TA mass (Figure 5a). In contrast, the GDF8 I56E and GDF8 H112A mutants reduced muscle mass by ∼25% (Figure 5a), which is consistent with the *in vitro* finding that these mutations enhance the ligand activity (Figure 3). Histological analysis, using hematoxylin and eosin staining of GDF8-treated TA muscles, revealed that the decreased muscle mass was a product of muscle fiber atrophy (Figure 5b), as indicated by decreased fiber diameter (Figure 5c). GDF8 I56E and GDF8 H112A also provoked a significant endomysial cellular infiltration, which was not evident in WT GDF8-treated muscles (Figure 5b). As we have shown previously with Activin A, these cells are likely collagen-secreting myofibroblasts (40) and their presence is indicative of enhanced GDF8 activity. Collectively, these data indicate that activating mutations in GDF8 markedly increase *in vivo* activity of this TGFβ superfamily ligand.

**Figure 5:**
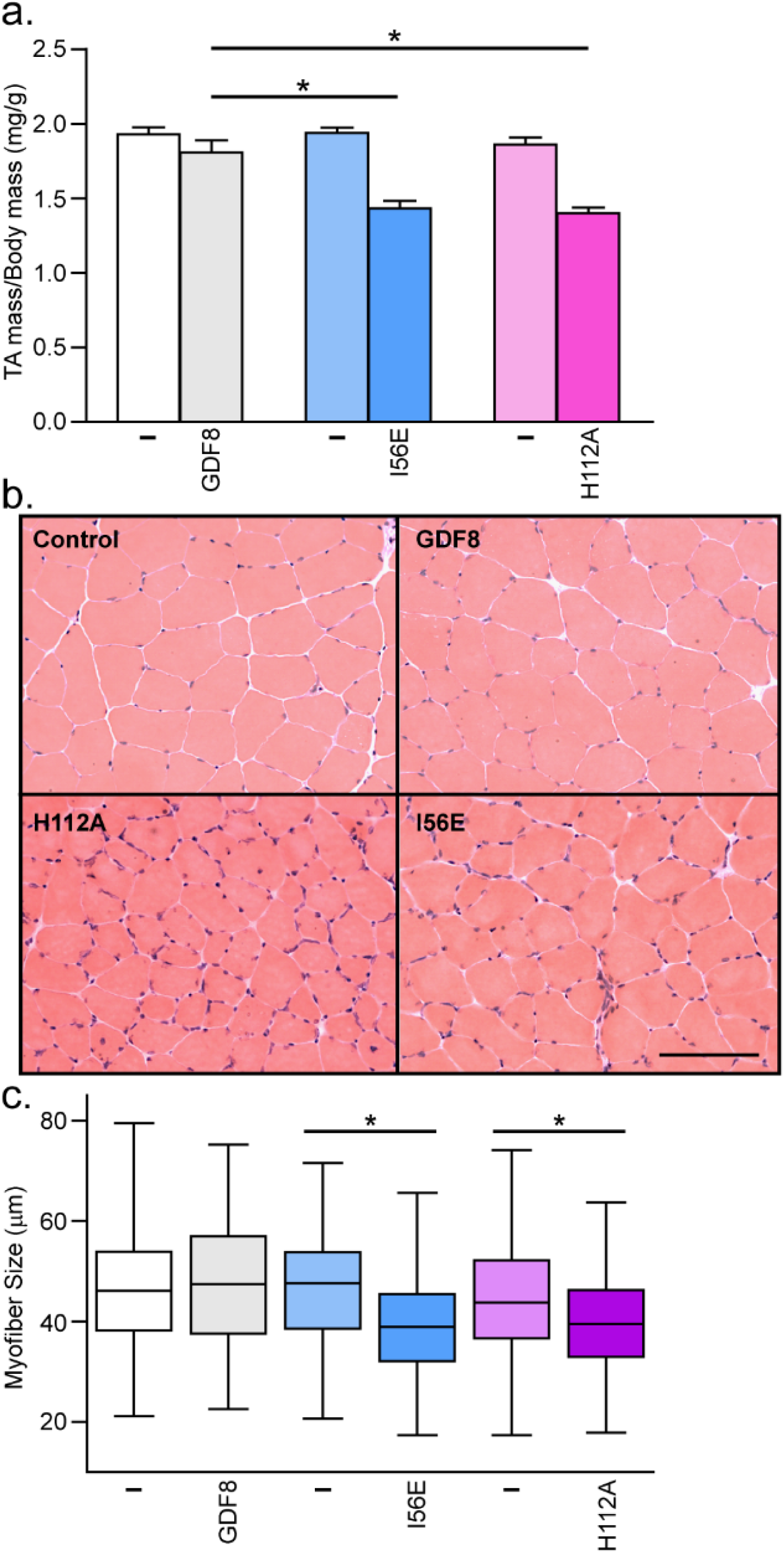
Activating mutations in GDF8 increase *in vivo* activity. The right tibialis anterior (TA) muscles of six-to-eight-week-old male C57Bl/6 mice were injected with AAV vectors encoding for GDF8, GDF8 (I56E) or GDF8 (H112A) (left TA muscles were injected with equivalent doses of an AAV vector lacking a transgene). **a)** Eight weeks post-AAV injection, the TA muscles were harvested and weighed (*n* = 4-6, paired Student’s t-test, data groups with different letters achieved significance, *p*<0.05; * significantly different to wild type GDF8, *p*<0.05). **b)** Hematoxylin and eosin staining of TA muscles was performed on cryosections (scale *bar* = 100 μm) and **c)** muscle fiber diameter quantified (*n* = 3, paired Student’s t-test, data groups with different letters achieved significance of *p*<0.05, at least 150 myofibers were counted per TA muscle).

## Discussion

The goal of this study was to elucidate mechanisms of latent GDF8 activation and identify the residues within the prodomain that contribute to latency. Although many ligands have been shown to loosely associate with their prodomains, only the prodomains of TGF-β ligands, GDF8, and the highly-related GDF11, have been shown to potently inhibit their respective ligands (reviewed in (14)). Through sequence alignment and structural modeling, we hypothesized that, despite the different modes of activation, that the GDF8 prodomain confers latency through a similar binding mechanism as observed in the latent TGF-β1 crystal structure (23). Using the low-resolution solution-based technique, SAXS, we demonstrated that the GDF8^L^ complex exhibits an ‘open’ conformation, unlike the ‘closed’ conformation adopted by the latent TGF-β1 structure. This difference is not unexpected given the mechanistic differences required for their respective activation. It is possible that an ‘open’ conformation is required for TLD activation of GDF8^L^ in order to improve accessibility of the TLD-cleavage site or TLD-recognition motif whereas a ‘closed’ conformation may impede access. However, through site directed mutagenesis of the GDF8 prodomain, based on sequence alignment to TGF-β1, we identified important residues within either the alpha-1 helix or fastener region, which when mutated, significantly enhance ligand signaling activity *in vitro* and *in vivo*. Together, our data supports the conclusion that the GDF8 and TGF-β1 prodomains both utilize similar residues to confer latency yet, we have identified that significant overall structural differences exist between the two complexes.

Apart from biological mechanisms of activation, it has been shown that exposure of latent TGF-β (25, 26) and GDF8^L^ (16, 31) to acidic conditions results in activation of the latent complexes. A molecular explanation for this mode of activation has yet to be determined but it has been postulated that ‘acid activation’ causes the prodomain and mature domain to dissociate, thus explaining the gain in ligand activity (16). Interestingly, our biophysical data strongly support that acid activation of the GDF8^L^ does not dissociate the complex but rather the pro- and mature domains remain associated, yet in a different molecular state, referred to as a ‘triggered’ state. Moreover, we determined that reconstitution of the GDF8 prodomain:ligand complex (GDF8^R^) from individual components did not result in a fully latent complex as the GDF8^R^ complex shows significant activity compared to the GDF8^L^ complex, suggesting that the latent state and ‘triggered’ state are not fully reversible. The notion that the GDF8 prodomain:ligand complex may exist in multiple activity states may explain, in part, why bacterially-derived and refolded GDF8 prodomain:ligand complex has been shown to have significant ligand activity (41) and, therefore may better represent the acid-activated or ‘triggered’ state. Nonetheless, our findings raise the possibility that mature GDF8 may be held in a locked or ‘spring-loaded’ state by its prodomain following biosynthesis, which can be ‘triggered’ when exposed to changes in pH.

In order to extend our understanding of the molecular interactions that drive GDF8 latency, we performed a targeted mutagenesis on the GDF8 prodomain, based on the latent TGF-β1 structure (23) and corresponding sequence alignment. Consistent with our hypothesis, we identified specific residues in the alpha-1 helix and the fastener region that when mutated, resulted in a more active ligand compared to WT whereas mutation of hydrophobic residues in the latency lasso region did not increase activity. Importantly, our data suggests that the increase in activity was not due to increased protein expression (Supplemental Figure 1). In fact, our most active mutant, I56E, showed the least detectable expression, perhaps due to rapid turnover of the mature ligand following receptor binding. Nonetheless, this observation is consistent with other groups that observed a reduction in ligand detection when corresponding residues were mutated in other TGF-β growth factors, though the effect of these mutations on TGF-β latency was not tested (34, 42).

Due to the overall complexity of GDF8 biosynthesis, latency, and activation, we are unable to define the molecular mechanisms to describe or explain why these mutations enhance GDF8 activity. Surprisingly, all activating GDF8 mutants required the presence of TLD except the I56E mutant, which remained significantly active despite the incorporation of the TLD cleavage-resistant mutation, D99A (GDF8 I56E/D99A; (24)). It is possible that incorporation of the I56E mutation disrupts the interaction between the alpha-1 helix and the mature ligand, which allows competition with GDF8 receptors. On the other hand, mutation of these regions may prevent GDF8 from fully entering the latent or ‘spring-loaded’ state during biosynthesis. Instead, this variant may be secreted in a form similar to the ‘triggered’ state that we have identified. This idea is supported by our data showing that the recombined mutant GDF8 prodomain:ligand complexes had similar signaling activity as GDF8^apo^. It is clear that further characterization of the mutant GDF8 prodomain:ligand complexes is necessary to pinpoint the molecular mechanism responsible for enhanced activity. However, we have identified specific residues within the GDF8 prodomain that can be modified to alleviate ligand latency without disrupting the function of the prodomain in folding and biosynthesis of the mature ligand (12, 13).

We extended our analysis of the GDF8 prodomain activating mutants *in vivo* using a model of skeletal muscle atrophy to determine if these mutants recapitulated our *in vitro* experiments. We focused our efforts on I56E, which showed the greatest activity independent of TLD and H112A where increased activity is completely dependent on TLD. In both cases, AAV delivery of I56E or H112A decreased the size of the muscle fibers relative to control mice and mice that received AAV encoding WT GDF8. Similar to our *in vitro* experiments, we were unable to detect evidence of mature GDF8 in the muscle of mice that received the AAV encoding the I56E mutant whereas we could reliably detect the mature ligand in the muscle from mice that received either the AAV encoding the H112A or WT proteins. As mentioned above, we speculate that loss of latency independent of TLD may enhance ligand turnover rate, thus making it challenging to detect the mature ligand. Together, these results are consistent with our previous observation that mutation of these residues results in a more active ligand.

While this manuscript was in preparation, it became apparent that an unpublished X-ray crystal structure had been determined in the laboratory of Marko Hyvonen (Figure 6; (43)). Therefore, we wanted to compare how our low-resolution SAXS data compared to the overall shape of the GDF8 prodomain:ligand complex. Consistent with our initial SAXS-based observations, the crystal structure of the GDF8 prodomain:ligand adopts a more open conformation that is drastically different from that of TGF-β1 and more similar to that of Activin A or BMP9. In agreement with our hypothesis that GDF8 prodomain contains similar inhibitory elements and mechanisms of TGF-β1, the alpha-1 helix, latency lasso, and fastener features are all present in the GDF8 prodomain:ligand complex. However, the conformation of the GDF8 prodomains in relation to their mature domain with which monomer they interact is significantly different than that of TGF-β1. For instance, the prodomain of one TGF-β1 monomer sits atop the other monomer of the homodimer, with all inhibitory elements imposed by one prodomain. However, the prodomain of one GDF8 monomer crosses over to interact with both mature domains of the dimer (Figure 6). Notably, the alpha-1 helix and latency lasso inhibit the GDF8 monomer from the same chain while the fastener interacts with the adjacent monomer. The significance of this binding strategy on inhibition is unknown. However, this ‘fastener-swap’ may play a role to ensure homodimer formation and/or aid in exposure of the TLD protease site. Nevertheless, I53 and I56 in the alpha-1 helix are shown to interact directly with the GDF8 ligand. It is possible that mutation of I53 or I56 would destabilize the alpha-1 helix and disrupt binding of the prodomain to GDF8. One would also expect that mutation of I60 would show a similar, if not more, exaggerated phenotype as the I53 or I56 mutants. However, mutation of I60 did not result in enhanced activity, but rather even lower activity than WT GDF8. It is possible that I60 may be important for protein folding and loss of this residue is detrimental to this process. Furthermore, the GDF8 prodomain:ligand crystal structure supports our finding that mutation of the fastener residues, Y111 and H112, would destabilize the fastener-interaction with the alpha-1 helix. This is similar to TGF-β1 where mutation of the fastener residues created a more active TGF-β1 ligand (23). Taken together, our mutational analysis of the GDF8 prodomain is highly consistent with the structure of the prodomain:GDF8 complex and also consistent with previous truncation analysis (41, 44-46). Our results are also consistent with unpublished results from the laboratory of Tim Springer who performed a rigorous hydrogen-deuterium exchange followed by MS to map the interactions of the prodomain with the mature in solution (47).

**Figure 6.**
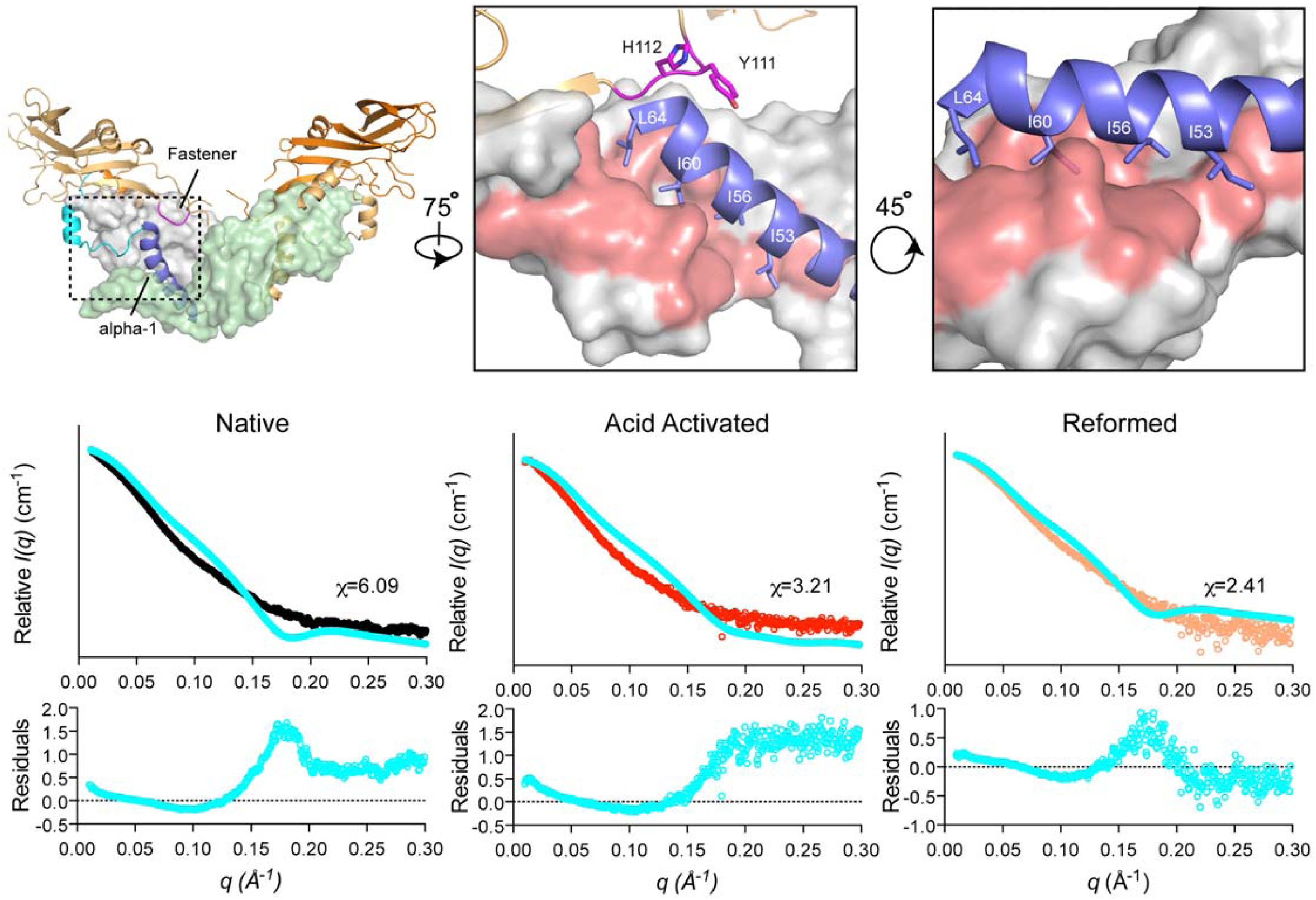
Structure of the GDF8 prodomain complex. The GDF8 prodomain complex containing the mature dimer (grey and pale green), alpha-1 helix (blue), latency lasso (cyan), fastener (magenta). Depiction of the alpha helix and fastener following a 75^o^ rotation about the y-axis (middle) residues. Sticks and labels highlight the residues I53, I56, I60, I64 within the alpha 1 helix, Y111 and H112, within the fastener. Right: depiction of the alpha-1 helix contacting the hydrophobic pocket (red) of the GDF8 monomer. The theoretical scattering profile derived from the GDF8 prodomain complex crystal structure compared to our experimental SAXS data on the various GDF8 prodomain complexes is shown at the bottom. The chi (X) value, determined by FoXS (32), for each comparison is shown adjacent to each scattering profile.

In summary, we determined that the latent GDF8 prodomain:ligand complex adopts a more ‘open’ structural conformation unlike that of the TGF-β1 latent complex (reviewed in (14)). Interestingly, both ligands share commonality with respect to the alpha-1 and latency lasso inhibitory elements, but show significant divergence with respect to the coordination of their respective fastener regions to confer latency. While it is unknown how this binding mode impacts or confers latency to GDF8 compared to TGF-β1, our data strongly supports the notion that the GDF8 prodomain:ligand complex can exist in multiple conformational states which ultimately dictate ligand activity and that the interactions between the prodomain and mature domain can be modified to generate a less latent and more active signaling ligand. It is plausible that GDF8 circulates within serum (17) in these various conformational ‘activity’ states, thus making it tempting to speculate that GDF8 biological regulation may include shifts in the balance of these ‘activity’ states depending on the physiological context.

## Materials and Methods

### HEK293-(CAGA)_*12*_ luciferase-reporter assay

Luciferase reporter assays for activation and inhibition were performed as previously described (27-31). Briefly, HEK293 (CAGA)_12_ cells (from RRID: CVCL_0045) stably transfected with plasmid containing Firefly luciferase reporter gene under the control of SMAD3-responsive promoter were seeded in growth media at 20,000 cells per well in a 96-well poly-D-lysine coated flat-bottom plate (Cat. No. 655940 Greiner Bio-One GmbH, Germany) and incubated at 37°C / 5% CO_2_ until 75-85% confluent. For transient expression experiments, 200 ng total DNA in a final volume of 25 μL (25-75 ng ligand DNA, 50 ng full length human furin in pcDNA4, 5-50 ng of appropriate TLD DNA in pRK5 or pcDNA3, filled to 200 ng with empty vector) per well was added directly to the growth media, incubated for 6 h and exchanged into serum-free media. OPTI-MEM reduced serum media (31985-070, Gibco, Life Technologies, USA) and TransIT-LT1 Reagent (MIR 2300, Mirus Bio LLC, USA) were utilized for transfection according to manufacturer instructions. Cells were lysed 30 h post-transfection using 20 μL per well 1x Passive Lysis Buffer (E1941, Promega, USA), on a plate shaker (800 rpm, 20 min, 20°C). The lysates were transferred to opaque black and white 96 well plates, 40 μL of LAR (E1501 and E1960, Promega, USA) was added, Firefly luminescence was recorded on Synergy H1 Hybrid Plate Reader (BioTek). When necessary, subsequent addition of 40 μL of Stop&Glo substrate (E1960, Promega, USA) was added and Renilla luminescence was recorded. To determine EC_50_ and IC_50_ values, the growth media was removed and the appropriate dilutions of either ligand alone or with antagonist, respectively, were serially titrated, and added to the cells in a 100 μL total volume of serum-free media. Luminescence was recorded as mentioned 18-24 h post ligand or antagonist addition. Experiments were independently performed at least 2 times and all data points were performed in triplicate. The EC_50_ and IC_50_ values were derived from non-linear regression with variable slope using GraphPad Prism 5 software. The EC_50_ and IC_50_ mean and standard error was calculated for each experiment and the mean weighted to the standard error was calculated using the following formulas, where ‘*a*’ is the standard error of the EC_50_ or IC_50_ determination, etc. (48):

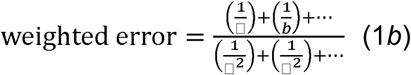

and

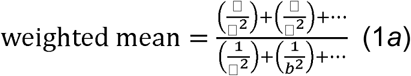

### Production and purification of GDF8 prodomain from E. coli

The prodomain of human GDF8 (residues 24-262) was cloned into a modified pET28a expression vector that contains an N-terminal 6x histidine tag, maltose binding protein (MBP) containing the mutations D82A/K83A/E172A/N173A/K239A for surface entropy reduction (49), and a HRV-3C protease cleavage site (6xHis-MBP-HRV3C cleavage site-GDF8 (residues 24-262)). The cysteine residues in the human GDF8 prodomain (C39/C41/C137/C138) were mutated to serine to improve expression and solubility and were shown to form a stable complex with mature GDF8 similar to mammalian derived GDF8 prodomain. *E.coli* Rosetta (DE3) strain carrying the appropriate prodomain construct was grown at 37°C, 220 rpm until an optical density (OD) of 0.8 at 600 nm was achieved, followed by cold induction with 0.5 mM IPTG, addition of 2% ethanol, and incubation at 20°C overnight. Cells were lysed and soluble 6xHis-MBP-GDF8 prodomain was applied to nickel affinity column (GE Lifesciences) equilibrated in 20 mM Tris pH 7.4, 500 mM NaCl followed by elution with a linear gradient using 20 mM Tris pH 7.4, 500 mM NaCl, 500 mM imidazole over 5 column volumes. The eluted protein was then dialyzed into 20 mM Tris pH 7.4, 500 mM NaCl and HRV-3C protease was added and incubated for 24 h to remove the 6xHis-MBP-fusion protein. Following cleavage, the protein was dialyzed into 10 mM HCl and applied to a C4 reverse phase column (Sepax) equilibrated in 0.1 % TFA, 5 % acetonitrile and eluted with a linear gradient to 0.1 % TFA, 95 % acetonitrile over 30 column volumes. The fractions containing GDF8 prodomain protein were pooled and buffer exchanged into 10 mM HCl for storage at −80° C for future use.

### Mammalian derived latent GDF8 complex (GDF8^*L*^)

Chinese hamster ovary (CHO) cells stably producing GDF8 were used as previously described (27, 30, 31, 50, 51). Conditioned media containing GDF8 was concentrated ∼10-fold using tangential flow and buffer exchanged into 50 mM Tris pH 7.4, 500 mM NaCl and applied to a Lentil Lectin-Sepharose 4B (Amersham Biosciences) column. Elution of GDF8 was conducted using the same buffer containing 500 mM methyl mannose followed by application to an S200 size exclusion column (Pharmacia Biotech; Buffer 20mM HEPES 7.4 500mM NaCl). Molarity of the GDF8 latent complex was determined as previously described, using SDS-PAGE/Coomassie staining and the quantified GDF8 mature as a standard (31).

### Acid Activation

Acid activation of GDF8 complex was performed as previously described (16, 31). In short, GDF8 complex was acidified to pH 2 - 7 using 1M HCl and incubating for 1 h followed by neutralization with 1M NaOH back to pH 8. Conversely, when a pH>8 was required 1M NaOH was used which was neutralized accordingly with 1M HCl. This material was then used in luciferase, and SAXS analysis.

### Small Angel X-ray Scattering (SAXS)

SAXS data was collected using SIBYLS mail-in SAXS service (Berkley, CA). GDF8 latent complex was purified as described above with the exception that the protein was reapplied to a Phenomenex HPLC S2000 size exclusion column equilibrated with 20 mM HEPES pH 7.4, 500 mM NaCl, 1 mM EDTA, 2 % glycerol. Generation of the reformed GDF8 complex required separation of mature GDF8 from the prodomain using previously described methods (27, 30, 31, 50, 51). Briefly, following purification of the GDF8^L^ complex, the complex was adjusted to 4 M guanidinium hydrochloride, 0.1 % TFA and applied to a C4 reverse phase column (Sepax). The fractions containing either the mature ligand or prodomain were then identified and quantified. The two proteins were then mixed together with an excess molar ratio of prodomain to mature ligand dimer (2.25 prodomain:1 ligand dimer) and neutralized with 100 mM HEPES pH 7.4, 500 mM NaCl. The protein was then applied to a Phenomenex HPLC S2000 size exclusion column as described above. Fractions from each peak were analyzed using SDS-PAGE followed by Western analysis to ensure that both proteins were present. Acid-activation of GDF8^L^ complex was performed as described above. Data were collected on purified at least 2 concentrations of GDF8^L^, GDF8^AA^, and GDF8^R^ in 20 mM HEPES pH 7.4, 500 mM NaCl, 1 mM EDTA, 2 % glycerol at 10°C. Four exposure times of 0.5 s, 1 s, 2 s, and 5 s were collected. Exposures exhibiting radiation damage were discarded. Buffer matched controls were used for buffer subtraction. ScÅtter (SIBYLS) and the ATSAS program suite (EMBL) were used for data analysis. Comparison of the experimental scattering profiles to known crystal structures was performed using the FoXS webserver (32). A*b initio* molecular envelopes were calculated using from the average of 23 independent DAMMIN (ATSAS, EMBL; (52)) runs using P2 symmetry, averaged using DAMAVER (ATSAS, EMBL; (53)), and filtered using DAMFILT (ATSAS, EMBL). SUPCOMB (ATSAS, EMBL; (54)) was used to superimpose the crystal structure of the latent GDF8 protein complex (43)

### Western analysis

To test protein expression following transfection, 500,000 HEK293 (CAGA)_12_ cells mentioned above (from RRID: CVCL_0045) were plated in a 6-well plate coated with poly-D lysine and incubated at 37^o^C until 75-85% confluency. A mixture of 625ng of GDF8 DNA, 1.25ug of Furin, and 3.125ug of pRK5 EV was used totaling 5ug of DNA, ∼25x the DNA used in a 96-well in order to closely mimic conditions within our luciferase assay. OPTI-MEM reduced serum media (31985-070, Gibco, Life Technologies, USA) and TransIT-LT1 Reagent (MIR 2300, Mirus Bio LLC, USA) were utilized for transfection according to manufacturer instructions. 12hrs post-transfection media was removed and replaced with serum-free media. 30hrs post transfection media was removed and concentrated 25x and run under reducing condition on an SDS-PAGE gel. Standard western protocols were utilized and the anti-GDF8 antibody from RnD Biosystems (AF788) was used as described by the manufacturer. Western blot was developed using the SuperSignal West Pico detection reagent (ThermoFisher) per manufacture instructions and detected using the C-DiGit blot scanner (LI-COR).

### Protein Thermal Shift

Protein thermal shift assays were conducted on a OneStep real-time PCR system (Applied Biosystems), run by the StepOne Software v2.3, as described by the manufacturer. In short, 1 μg of protein was placed in 20 μl of 20 mM HEPES pH7.4, 500 mM NaCl in the presence of 1x ROX reagent from the Protein Thermal Shift Dye Kit^™^ (Applied Biosystems). The melting temperature and Tm of each protein was conducted on a 1% gradient from 25°C-100°C taking approximately 40 min. Data was analyzed using Protein thermal shift software v1.3, and curves were plotted from triplicate measurements using GraphPad Prism5 software.

### Production of AAV vectors

The cDNA constructs encoding for WT GDF8, GDF8 I56E and GDF8 H112A were cloned into an AAV expression plasmid consisting of a CMV promoter/enhancer and SV40 poly-A region flanked by AAV2 terminal repeats. These AAV plasmids were co-transfected with pDGM6 packaging plasmid into HEK293 cells to generate type-6 pseudotyped viral vectors. Briefly, HEK293 cells were seeded onto culture plates for 8-16 h prior to transfection. Plates were transfected with a vector-genome-containing plasmid and the packaging/helper plasmid pDGM6 by calcium phosphate precipitation. After 72 h, the media and cells were collected and subjected to three cycles of freeze-thaw followed by 0.22 μm clarification (Millipore). Vectors were purified from the clarified lysate by affinity chromatography using heparin columns (HiTrap^TM^, GE Healthcare), the eluent was ultra-centrifuged overnight, and the vector-enriched pellet was re-suspended in sterile physiological Ringer’s solution. The purified vector preparations were quantified with a customized sequence-specific quantitative PCR-based reaction (Life Technologies).

### Administration of AAV vectors to mice

All experiments were conducted in accordance with the relevant code of practice for the care and use of animals for scientific purposes (National Health & Medical Council of Australia, 2016). Vectors carrying transgenes of GDF8 variants were injected into the right tibialis anterior (TA) muscle of 6-8-week-old male C57Bl/6 mice under isoflurane anesthesia at 10^10^ vector genomes (vg). As controls, the left TA muscles were injected with AAVs carrying an empty vector at equivalent doses. At the experimental endpoint, mice were humanely euthanized via cervical dislocation, and TA muscles were excised rapidly and weighed before subsequent processing.

### Histological analysis

Harvested muscles were placed in OCT cryoprotectant and frozen in liquid nitrogen-cooled isopentane. The frozen samples were cryosectioned at 10 μm thickness and stained with hematoxylin and eosin or Masson’s Trichrome. All sections were mounted using DePeX mounting medium (VWR, Leicestershire, England) and imaged at room temperature using a U-TV1X-2 camera mounted to an IX71 microscope, and an Olympus PlanC 10X/0.25 objective lens. DP2-BSW acquisition software (Olympus) was used to acquire images. The minimum Feret’s diameter of muscle fibers was determined using ImageJ software (US National Institutes of Health, Bethesda, MD, USA) by measuring at least 150 fibers per mouse muscle.

## Acknowledgements

This work was supported, in part, by National Institutes of Health, the National Health and Medical Research Council Grant, the Muscular Dystrophy Association, the University of Cincinnati Graduate Dean Fellowship, and the American Heart Association (R01AG047131, R01AG040019, and R03AG049657 to RTL; 1078907 to CAH; App 1117835 for Gregorevic Senior Research Fellowship from NHMRC (Aust.); Graduate Dean Fellowship and 12PRE11790027 to RGW; R01GM114640 and Muscular Dystrophy Association Grant 240087 to TBT). This work was also supported by the Integrated Diffraction Analysis Technologies Program of the Department of Energy Office of Basic Energy Sciences awarded to the Advanced Light Source at Lawrence Berkeley National Laboratory. The Baker Heart and Diabetes Institute is supported in part by the Operational Infrastructure Support Program of the Victorian Government. The Authors thank Dr. Hongwei Qian (Baker Heart and Diabetes Institute) for assistance with production of recombinant AAV vectors.

### Author Contributions

RGW, TBT, and CAH conceived the idea and designed the experiments. RGW and JCM prepared all figures and wrote the initial draft of the manuscript, performed the experiments, purified and expressed the proteins required for experimentation. RGW identified the initial mutants presented in the manuscript and designed the initial HEK293 (CAGA)_12_ cell-based assays. JCM and MC performed the HEK293 (CAGA)_12_ cell-based assays. JCM designed and performed the thermal shift assays. MJM and RTL aided in the production and purification proteins. TC and MH solved the structure of the GDF8 prodomain complex. PG and CAH designed and performed the AAV experiments. All authors revised and edited the manuscript. All authors read and approved the final manuscript.

### Conflicts of interest

TBT is a consultant for Acceleron Pharma. The University of Cincinnati and Monash University have filed for intellectual property on GDF8 and GDF11 listing RGW, TBT, and CAH as inventors. Harvard University and Brigham and Women’s Hospital have filed for intellectual property on GDF11 listing RTL as an inventor. The other authors report no competing interests.

## Supplemental Figure Legends

**Supplemental Figure 1.**
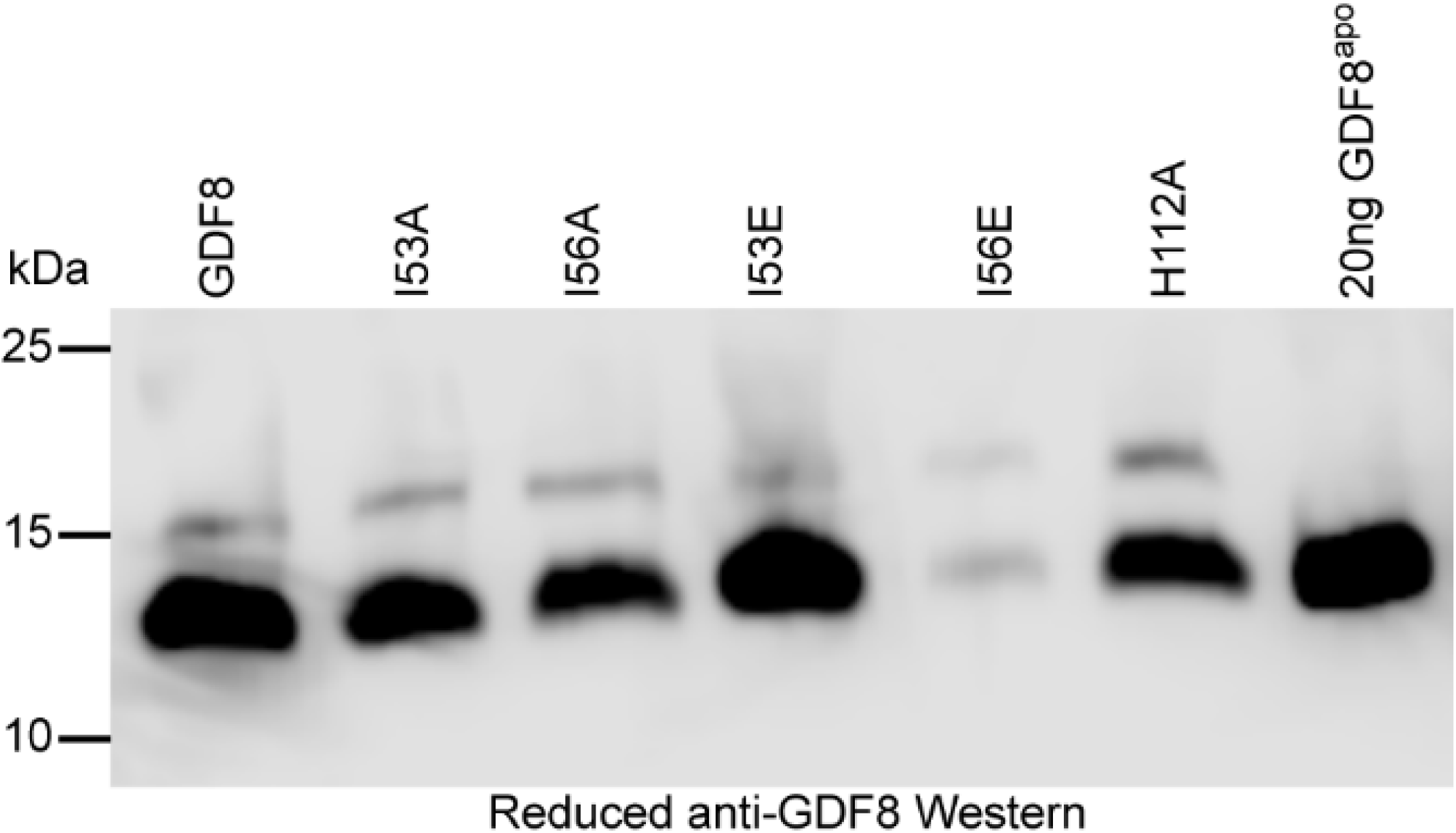
Relative protein expression following transient transfection of various GDF8 prodomain mutant constructs. Conditioned media from HEK293 (CAGA)_12_ luciferase cells probed for mature GDF8 under reducing conditions.

